# Automatic Generation of Interactive Multidimensional Phase Portraits

**DOI:** 10.1101/2022.02.23.481676

**Authors:** Oluwateniayo O. Ogunsan, Daniel Lobo

## Abstract

Mathematical models formally and precisely represent biological mechanisms with complex dynamics. To understand the possible behaviors of such systems, phase portrait diagrams can be used to visualize their overall global dynamics across a domain. However, producing these phase portrait diagrams is a laborious process reserved to mathematical experts. Here, we developed a computational methodology to automatically generate phase portrait diagrams to study biological dynamical systems based on ordinary differential equations. The method only needs as input the variables and equations describing a multidimensional biological system and it automatically outputs for each pair of dependent variables a complete phase portrait diagram, including the critical points and their stability, the nullclines of the system, and a vector space of the trajectories. Crucially, the portraits generated are interactive, and the user can move the visualized planar slice, change parameters with sliders, and add trajectories in the phase and time domains, after which the diagrams are updated in real time. The method is available as a user-friendly graphical interface or can be accessed programmatically with a *Mathematica* package. The generated portraits and particular views can be saved as computable notebooks preserving the interactive functionality, an approach that can be adopted for reproducible science and interactive pedagogical materials. The method, code, and examples are freely-available at https://lobolab.umbc.edu/autoportrait.

## 1. Introduction

Understanding and predicting the dynamics of complex biological systems require mathematical models able to abstract the specific interactions between their components [1]. Among them, dynamical systems based on differential equations are one of the most popular methods to formally and precisely represent biological mechanisms [2]. Crucially, multiple user-friendly tools are available for their formulation, numerical simulation, and sharing [3–7], which has contributed to their widespread use in biological research and education [8]. In addition to simulating their dynamics through time, visually analyzing the global behavior of such models is essential for understanding their function. However, the visual analysis of dynamical systems remains a challenging process in need of user-friendly tools.

Phase portrait diagrams are ideal for visualizing the overall behavior of a biological dynamical system [9,10]. Such portraits can include a vector field of the gradient and direction of the trajectories in a domain, nullclines representing horizontal or vertical trajectories, and critical points representing the equilibrium states in the system [11]. Complete phase portraits are generally limited to present information in two dimensions. However, dynamical systems with more than two dependent variables can be visualized without loss of information with state combinations [12]. In general, constructing phase portrait diagrams requires expert ability and the use of specialized mathematical software tools.

Despite the numerous user-friendly tools for the time-course simulation of biological dynamical systems [13], the generation of phase portraits remains a highly-specialized task without fully-automated tools. General-purpose packages for mathematical analysis such as MATLAB, MAPLE, and *Mathematica* are able to produce phase portraits, but require the implementation of custom scripts and knowledge of the particular mathematical libraries and plotting functions in these packages [14,15]. Specialized software tools have been developed for the analysis of dynamical systems with the ability to generate phase portraits. *XPPAUT* is a popular and powerful application for simulating, animating, and analyzing dynamical systems [16], with a particular focus on the efficient numerical solving of differential equations. *XPPAUT* can be used to generate phase portraits, although every element and plane need to be added to the plot by the user every time a system is opened with the tool. *ASysViewer* is an application for the interactive visual exploration of autonomous dynamical systems using line integral convolution techniques [17], but it is restricted to two dimensional systems, limiting its applicability. Another limitation of these tools is their inability for sharing a specific diagram that could be further interacted with and analyzed in the form of a dynamical publication or pedagogical material. Such a tool could be used in research manuscripts to include interactive results and benefit reproducible science and education in dynamical systems and systems biology.

Here, we present a novel tool called *AutoPortrait* for the automatic generation of interactive phase portraits for multi-dimensional ordinary differential equations. The tool can be used both with a user-friendly interface for directly typing the equations and with a *Mathematica* package to be called from scripts. Once a phase portrait is generated, it can be embedded into a notebook saving the exact view while keeping all the interactive functionality. The tool automatically detects the parameters in the equations and then generates a two-dimensional phase portrait for every combination of two dependent variables, choosing an appropriate value for the other free variables, parameters, and domain range. The tool computes and displays in the portrait a vector field of the trajectories, the nullclines of the system, and the critical points with their stability. A user can interact with the generated diagrams using the mouse, which includes panning and zooming the plot, moving sliders to select a different plane, clicking on critical points to center them, and visualizing trajectories in the phase and time domain. We include several examples illustrating the use of the tool to understand biological systems.

## 2. Computation of multidimensional phase portraits

We outline the automatic method to compute complete phase portrait diagrams including critical points, vector fields, and nullclines. Portraits including all these features are best visualized in two-dimensional plots and the method produces all possible 2-dimensional views in phase space.

We consider an *n*-dimensional autonomous dynamical system

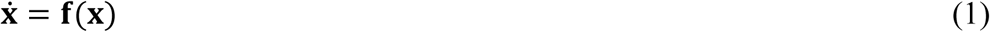

where **x** = (*x*_1_, …, *x*_*n*_) are variables in an open connected subset Ω ⊂ ℝ^*n*^, an overdot denotes differentiation with respect to time, *t*, and **f**(**x**) = *f*_1_(**x**), …, *f*_*n*_(**x**) are continuously differentiable functions that only depend on the variables. Thus, the variables might represent any biological entity modeled in a continuous fashion, such as chemical concentrations, gene expression, or large populations of different species. The functions might represent their production or growth rates, such as chemical reactions, gene regulation, or species competition, respectively.

The existence and uniqueness theorem ensures that starting from any point in Ω, a solution exists for the system of equations (1) and that the trajectories formed by all solutions never intersect. Hence, we can consider the system (1) as a *n*-dimensional vector field with a vector 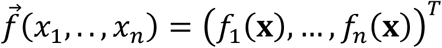 attached to each point (*x*_1_, …, *x*_*n*_) ∈ Ω. A stream plot is an intuitive graphical representation of a vector field, but the dynamics can only be completely visualized in 2-dimensions. Hence, multiple plots will be necessary to understand systems with more than two dependent variables.

Phase portraits are defined in phase space, where the axes of the plot represent dependent variables instead of independent variables such as time. For an *n*-dimensional system, the problem arises in how to describe the dynamics of the system reliably. From an *n*-dimensional system, a 2-dimensional phase plot can be generated representing a planar slice showing the dynamics of the system with respect to only two dependent variables and setting the rest of dependent variables as constants. Crucially, the location of a *n*-dimensional point or line in Ω can be unequivocally determined from all the combinations of 2-dimensional plots of *x*_*i*_, *x*_*j*_ out of the set (*x*_1_, …, *x*_*n*_) [12]. Hence, a number *v* of 2-dimensional views equal to the binomial coefficient 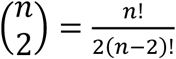 is sufficient to determine unambiguously the location of critical points, nullclines, and the numerical solution of a trajectory in an *n*-dimensional system. In this way, 2-dimensional systems require one view (*x*_1_, *x*_2_), 3-dimensional systems require three views (*x*_1_, *x*_2_), (*x*_1_, *x*_3_), and (*x*_2_, *x*_3_), 4-dimensional systems require six views, and so forth. This approach has been used before to visualize the trajectories in the phase plane of three-dimensional systems [18,19].

In this manner, the method generates 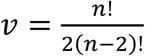 phase portraits to describe the phase portrait elements of an *n*-dimensional system. Each phase portrait *p* ∈ (1, …, *v*) represents a plane slice of the system through variables (*x*_*i*_, *x*_*j*_), and hence the rest of the variables *k* ∈ (1, …, *n*)\(*i, j*) are set as constants 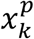 such that 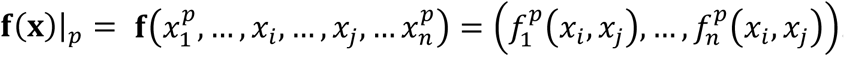. Each of the phase portraits also can include nullclines, which indicate where the trajectories are purely horizontal or vertical. For a phase portrait *p* of (*x*_*i*_, *x*_*j*_), the nullclines are defined as the curves in which 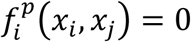 or 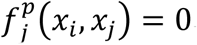.

In addition to the vector fields and nullclines showing the general dynamics of the system, critical points (also called fixed or equilibrium points) and their stability can identify states of equilibrium within the system. A critical point **x**^*^ is defined by **f**(**x**^*^) = **0**, and its stability is characterized by linearizing the system (1) and analyzing the eigenvalues of the Jacobian matrix

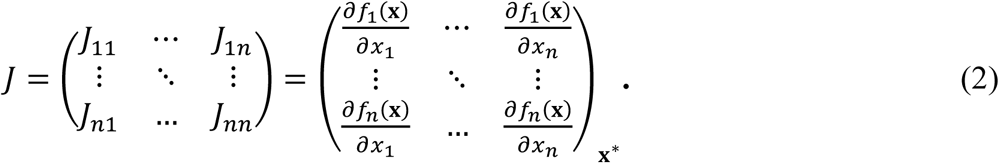

A critical point is classified as stable if all eigenvalues have negative real parts, as unstable if all eigenvalues have positive real parts, as saddle if some eigenvalues have negative and others positive real parts, and as higher-order if at least one eigenvalue is zero. In the latter case, the critical point is nonhyperbolic, and although its stability cannot be obtained through linearization, it could be deduced visually from the stream plots.

Finally, the range of the phase portraits is selected automatically from the location of the critical points to ensure they are included in the visualized domain. Biological systems are often restricted to the nonnegative domain, since the dependent variables usually represent species populations, molecular concentrations, etc. In this way, the computation and visualized range of the phase portrait can be restricted to solutions in the nonnegative domain.

## 3. GUI and script implementation

The automatic methodology for creating phase portraits, called *AutoPortrait*, has been implemented in *Mathematica*, an ideal environment for modeling dynamical systems [15]. The method uses the solver *NSolve*, which automatically can select from several efficient numerical approaches, as well as the *Eigenvalues* function to find exact or numerical eigenvalues. Two different implementations of the method have been developed: (1) an interactive graphical user interface without the need of any programming knowledge and (2) a package that can be imported into any Mathematica notebook or code. The generation of portraits for new systems of equations requires the full version of *Mathematica*, but the interactive portraits generated with the package can be integrated into *Mathematica* notebooks and computational documents that can be opened with the freely-available *Wolfram Player*, preserving all the interactive functionality of the plots.

To first illustrate the basic use of the method interface, we define a simple Lotka-Volterra model of competition for food between two species [11] as

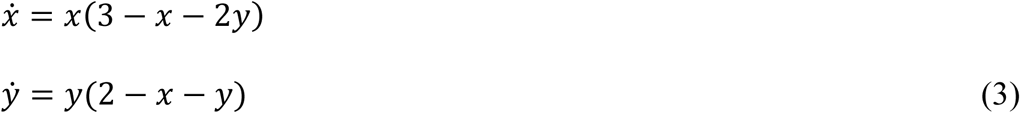

where *x* are a population of rabbits and *y* are a population of sheep. Both populations follow logistic growth, with a faster rate for rabbits, and a reduced growth due to conflict through food sharing proportional to the population sizes, with a larger effect on rabbits.

Figure 1 shows the graphical user interface of the method, where the equations for the derivatives (3) have been directly typed in the application and the resultant automatically-generated phase portrait is shown. A slider allows the user to select the number of dependent variables in the dynamical system, from two to 10, and a checkbox can be clicked to restrict the domain to nonnegative values. A system of two equations will generate a single phase portrait, while a system with 10 equations will generate 45 phase portraits (all possible combinations of two variables). The dependent variable names can be changed just by typing a new name in their box. Similarly, the equations for their derivatives can be typed in their respective input boxes. After the system is defined, the user can click the button *Generate portraits* after which all the interactive phase portraits will be presented on the screen, as shown in the figure. The phase portrait includes the stream plot, nullclines, and critical points and their stability. The method automatically computes an appropriate range that includes all the critical points. Finally, a legend with the symbols and colors used in the portraits as well as the mouse commands to interact with the portraits are displayed in a legend next to the generated plots. In this way, the graphical user interface allows any user with no programming knowledge to generate automatically detailed phase portraits.

**Figure 1.**
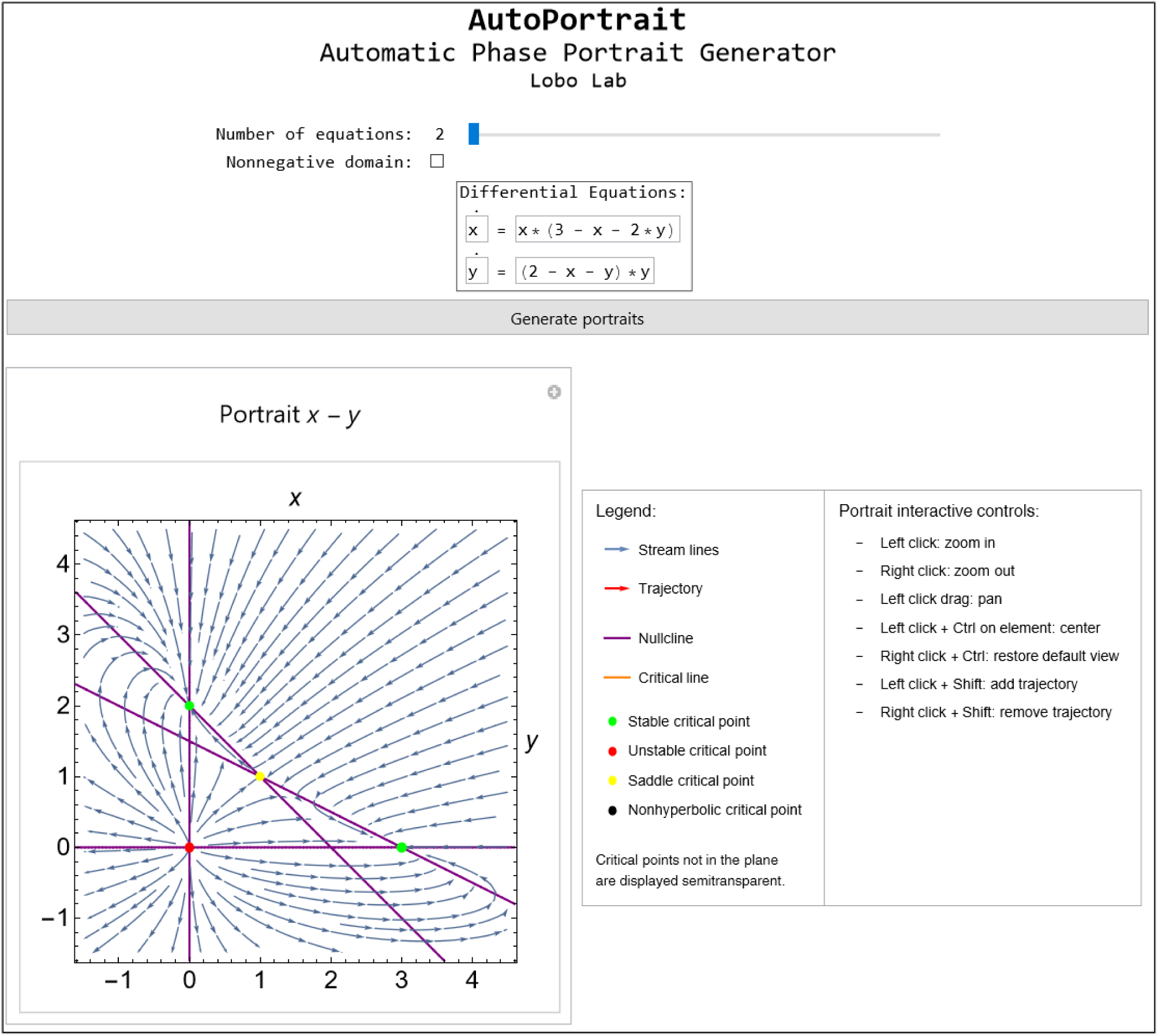
Main view of the user-friendly graphical user interface for the presented method showing the automatically-generated phase portrait for a Lotka-Volterra model of competition. The user can select the number of dependent variables and directly type their equations in the input boxes, after which the interactive phase portrait and legend are displayed.

Commonly, dynamical systems are studied on the basis of some parameters that modulate the dynamics of the system, a task that is facilitated with the presented methodology. Parameters typed in the equation boxes are automatically recognized and displayed next to the equations, where the user can select their initial value and range. When generating the phase portraits, a new set of input boxes and sliders are presented to interactively change the value of each parameter. The phase portraits are recomputed dynamically in real time as the user changes any parameter by typing in their respective input box or moving the slider. Figure 2 shows the interactive interface when typing the same Lotka-Volterra model of competition as in (3) but now including free parameters as follows

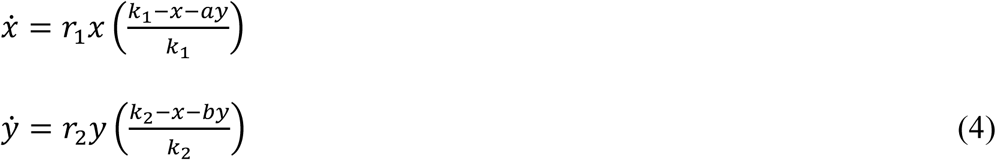

where *x* and *y* are the two populations, *r*_1_ and *r*_2_ are the intrinsic growth rates, *k*_1_ and *k*_2_ are the carrying capacities, and *a* and *b* are the competition coefficients. The parameter values chosen now changes the dynamics of the system so the two populations can coexist in equilibrium (green dot) [20]. Parameters can be easily changed interactively using the sliders or input boxes to observe their effect in the system dynamics in real time. Figure 3 shows an example of a bifurcation after changing the parameter *b* with the slider, which causes the nullclines to avoid an intersection in the positive domain and hence preventing the two populations to coexist.

**Figure 2.**
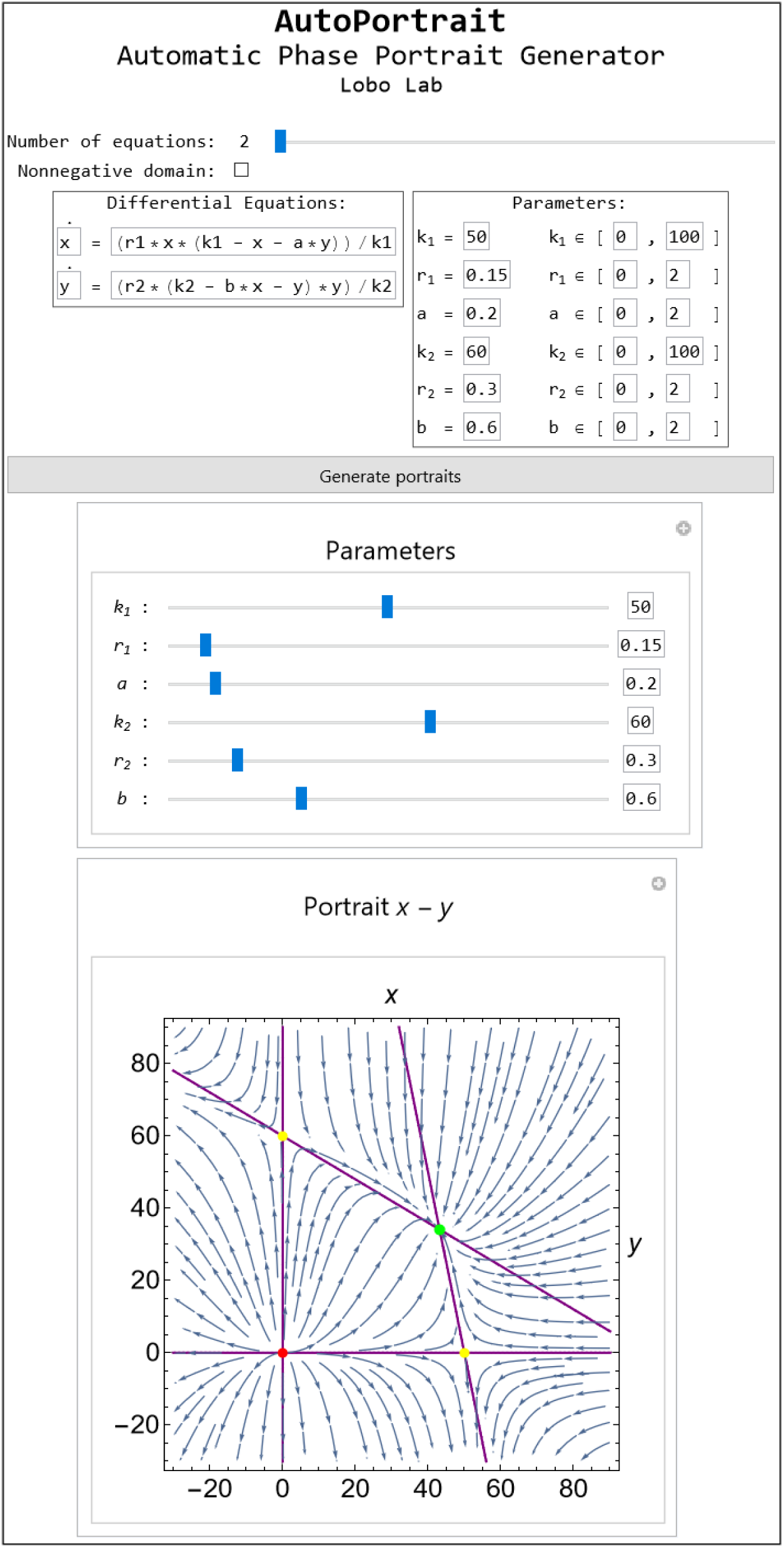
Main view of the graphical user interface and resultant phase portrait for a Lotka-Volterra model of competition with parameters. The method automatically recognizes the free parameters in the equations and present input boxes to select their initial value and range. After generating the portrait, a slider for each parameter is included above the portrait for the user to interactively change them and visualize their effect in the phase portrait in real time. Legend is omitted for brevity.

**Figure 3.**
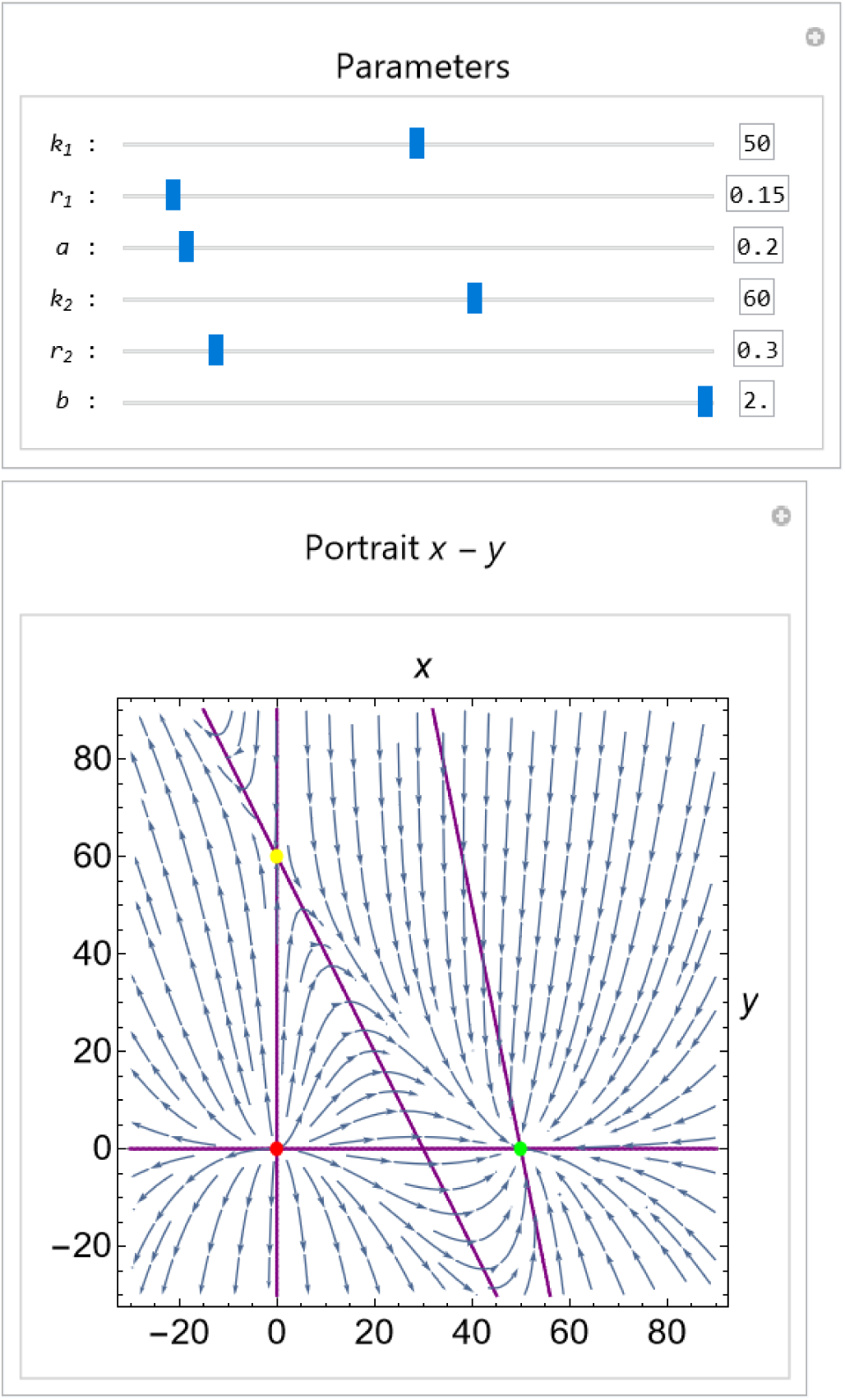
The generated portraits include sliders and input boxes for the free parameters in the equations. In this way, the user can change the parameter values and observe in real time their effects in the system dynamics. The plot shows how increasing the competition coefficient *b* in the Lotka-Volterra model of competition causes the extinction of one species in the stable equilibrium state.

In addition to the graphical user interface, the method is available directly as a *Mathematica* package that can be imported into any notebook or code. In this way, the method can be integrated with more complex scripts, such as exploring portraits for a family of equations. Box 1 shows the necessary simple commands to generate the phase portraits for the system (3), resulting in the same interactive phase portrait and legend as in Figure 1. The function *GeneratePortrait* requires two arguments. The first argument is the list of differential equations in the system. Notice that the independent variable for time is not necessary to be specified, which makes the function simpler to use. The second argument is a list of dependent variables. Similar to the interactive user interface, the method automatically recognizes the parameters in the equations and presents along the phase portraits sliders and input boxes when appropriate for the user to interact with them, as shown in Figure 2. In addition, the function includes optional arguments to set the initial parameter values, their ranges for the sliders, restrict the domain to nonnegative values, or include specific trajectories in the plot.

### 3.1. Interactive features

#### Box 1.

Mathematica script to automatically generate the phase portrait for equations (3) with the *AutoPortrait* package.

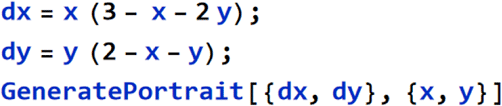

The phase portraits generated by both the graphical user interface and the package function are interactive to facilitate the exploration and understanding of complex dynamical systems (see the legend in Figure 1 for a list of interactive controls available). Hovering the mouse pointer on a critical point or nullcline displays its coordinates or line function, respectively. For systems with more than two dimensions (see next section), the values of the dependent variables not represented in the plot can be assigned with a slider or input box, effectively changing in real time the plane at which the phase portrait is shown. In addition, critical points can be clicked to move the plane to the location of that critical point. Crucially, the user interactively can change the range of the axes by dragging or clicking with the mouse anywhere inside a plot, which allows for panning and zooming in and out the plot.

Trajectories in the phase and time domains can be added and visualized in the plots. The user can select any point within a phase plot to display the projection of a trajectory starting in that point. This feature is especially useful for understanding the dynamics of systems with more than two dimensions, as all the portraits will include the projection of the same trajectory in their displayed plane. Figure 4 shows the same phase portrait as Figure 2 but restricted to the nonnegative domain and including a trajectory (red line) converging to the stable equilibrium state (green dot). Alongside the phase portraits, the same trajectory is computed and presented in the time domain. In this way, a user interactively can generate particular portraits emphasizing specific locations and coordinated trajectories in the phase and time domains.

**Figure 4.**
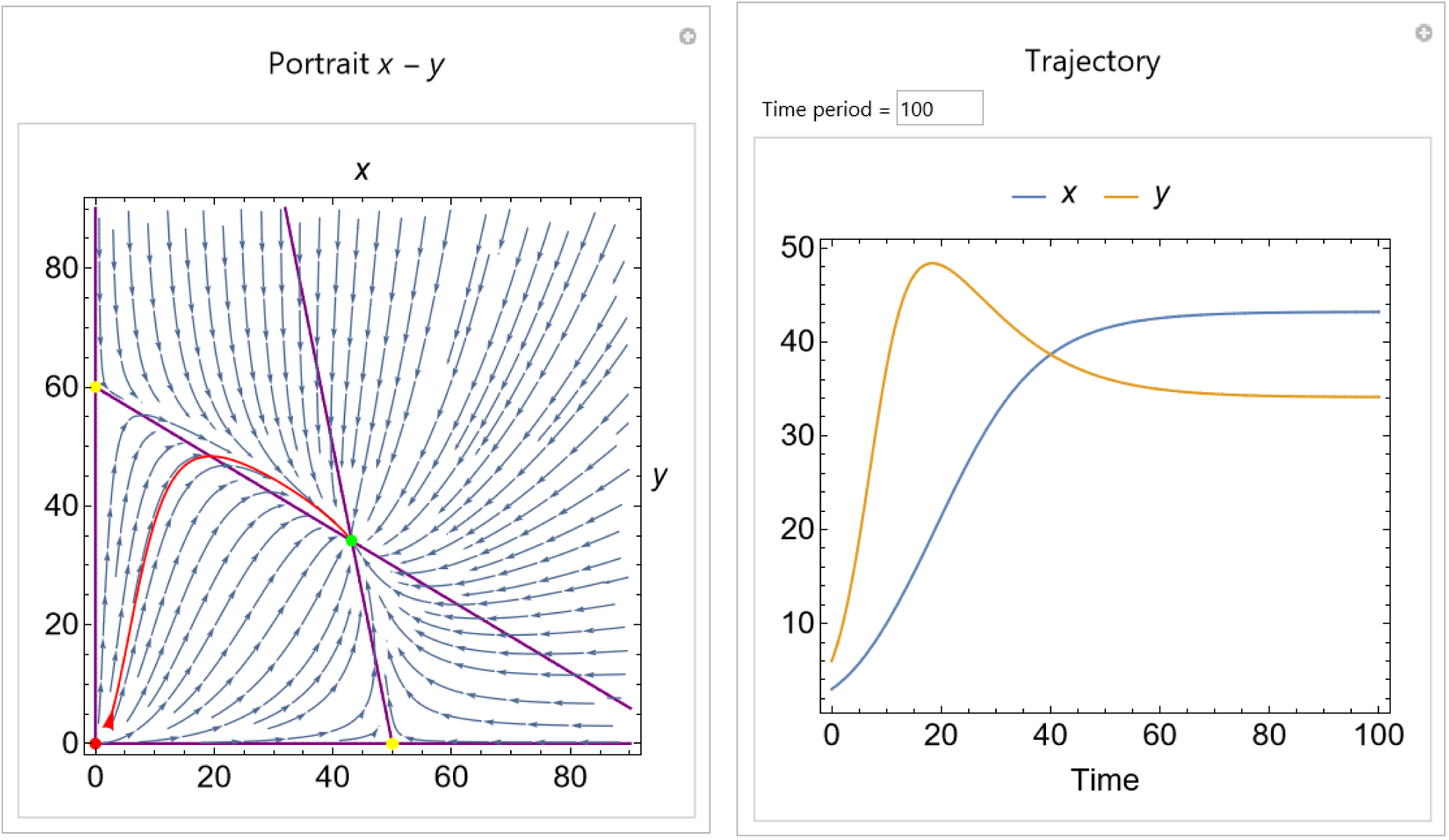
Phase portrait of the Lotka-Volterra model of competition generated by the method including a trajectory in the phase (red line) and time domains. The user can click on the phase portraits to generate a trajectory starting in that point. The initial point of the trajectory is shown with a red arrowhead.

Importantly, the interactive portraits generated by the methodology can be copied and pasted directly into other *Mathematica* notebooks and computable documents, retaining their interactive functionality. The complete interface can be saved, including a particular system, parameter values, plots, and trajectories, and also retaining the ability to be interactive when reopened. The notebooks including the plots can be opened with the freely-available *Wolfram Player*, allowing any user to interact with the system. This flexibility makes *AutoPortrait* an ideal tool for reporting portraits in dynamic publication supplements as well as for incorporating them in pedagogical materials for students to interact and understand the mechanisms of biological systems.

## 4. Higher dimensional systems

The presented method is especially useful for interactively understanding dynamical systems with more than two dimensions. This section will demonstrate this ability by using the method to generate phase portraits of a three-dimensional predator-prey model with infected prey and a four-dimensional Lotka-Volterra model of competition.

### 4.1. Predator-prey model with infected prey

Extending the classical Lotka-Volterra competition model of two populations with further species allows for the study of complex dynamics from social behavior interactions. We consider the three-dimensional ecoepidemic model of predator-prey interactions presented in [21], where the prey population exhibits group defense and are also affected by a transmissible disease. The system includes a population of healthy prey that grows logistically but is perturbed by both a population of predators and a spreading infectious disease modeled with a bilinear term. The diseased prey forms a third population that does not reproduce or contribute to density pressure but that can be hunted by the predator easier than the healthy prey as well as spread the disease to the healthy prey. In this way, the model is defined in its non-dimensionalized form with

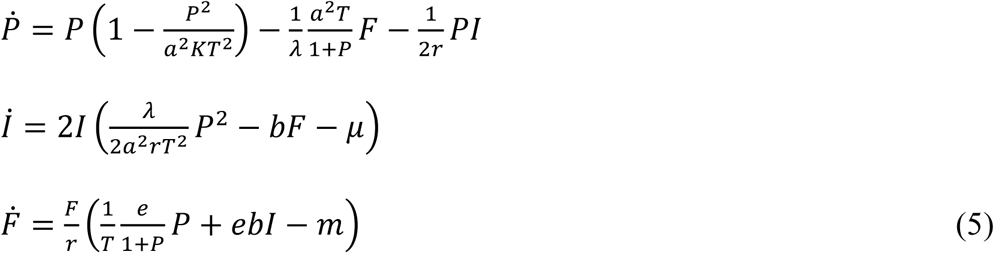

where *P* are the healthy prey population (squared due to non-dimensionalization), *P* are the infected prey population, *F* are the predator population, and the parameters are defined as in Table 1.

**Table 1.** Parameters for the predator-prey model with infected prey.

| Symbol | Value | Description |
| --- | --- | --- |
| $a$ | 0.5 | Hunting rate of predator on healthy prey |
| $e$ | 0.5 | Conversion factor of prey into predators |
| $K$ | variable | Carrying capacity of prey |
| $m$ | 0.42 | Natural mortality rate of predators |
| $r$ | 0.7 | Intrinsic growth rate of healthy prey |
| $T$ | 0.8 | Average time to capture a healthy prey |
| $\lambda$ | 0.7 | Infection rate |
| $b$ | 0.7 | Contact rate of predator and diseased prey |
| $\mu$ | 0.65 | Natural and disease mortality of infected prey |

The system presents a number of bifurcations [21] and here we employ automatically-generated phase portraits and trajectories with *Autoportrait* to illustrate the effect of the parameter *K* for the carrying capacity of prey. Figure 5 shows the generated phase portraits of the system at low prey carrying capacities. Three portraits are produced for each parameter value, each showing a 2D phase portrait with the vector plot at a particular plane for the third variable. The user can change the viewed plane independently with the slider above each phase portrait, which updates in real time the portrait. By default, the portraits are centered in the critical point with the least zero values (usually the most interesting in the system), but each of them can be clicked in the interface to center the view on their plane. A trajectory can be added to the phase portraits by either clicking the portrait or including a particular argument in the package function call. When clicking in a portrait, the initial value for the trajectory is updated according to the two dependent variable values in the portrait. To select the value for the other dimensions, the user can click the other portraits. The selected trajectory is projected across all the portraits, together with a new plot in the time domain visualized on the right. Through the interactive interface to select the viewed phase plane and plot trajectories, a user can easily understand the temporal dynamics of complex multidimensional systems.

**Figure 5.**
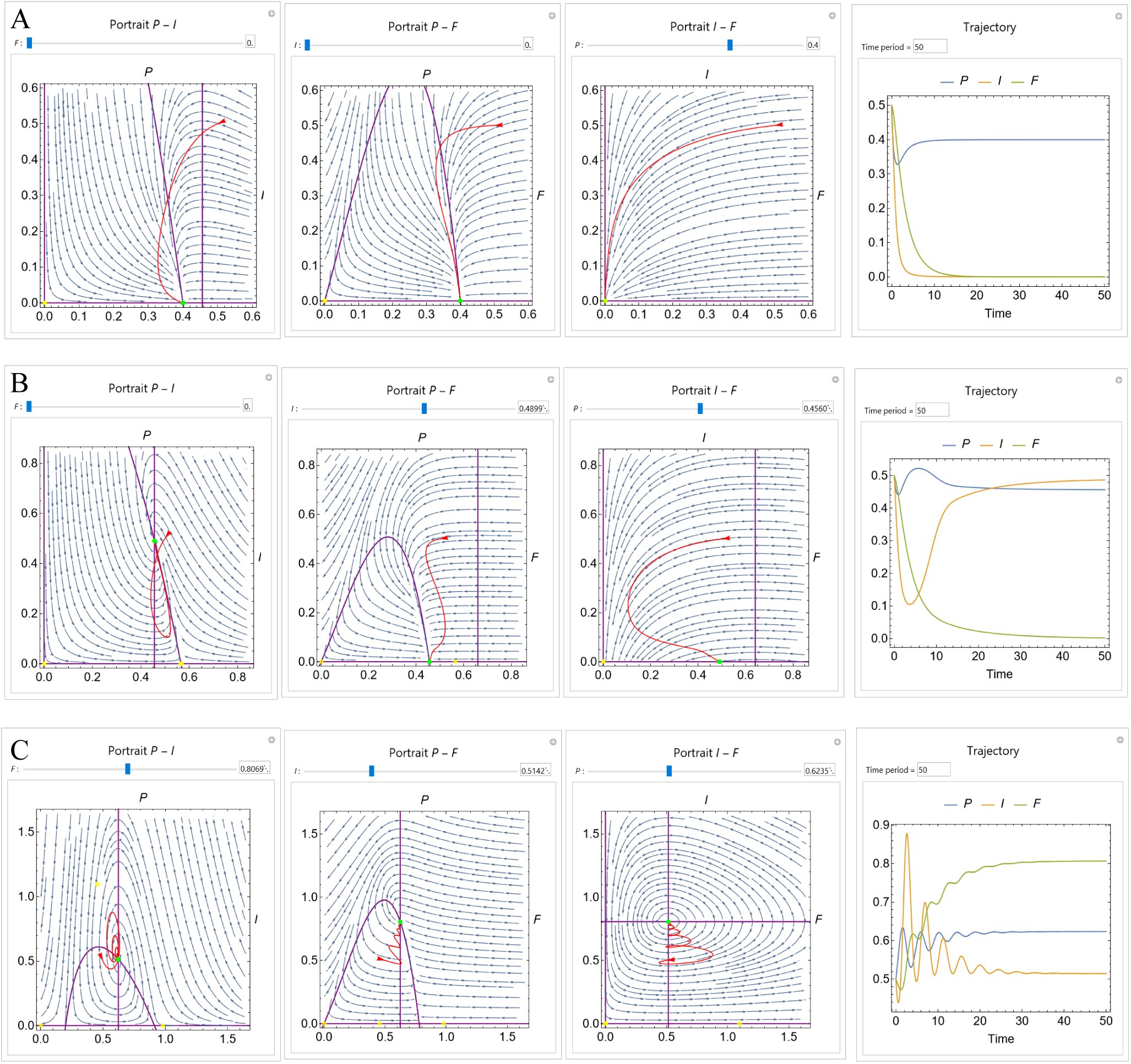
Automatically-generated outputs of the method for a predator (*F*), prey (*P*) model including prey group defense and an infected prey population (*P*) with low levels of prey carrying capacity *K*. All other parameters are set as in Table 1. All the portraits are automatically centered on the stable equilibrium point and show a trajectory (red and right panels) starting at a population of 0.5 for all the species. **A**. At low prey carrying capacity (*K* = 1), only the healthy prey population can survive. **B**. Increasing the prey carrying capacity (*K* = 2) through a transcritical bifurcation clearly visible in the *P* − *P* portrait, the infected prey population can survive together with the healthy prey. **C**. Further increasing the prey carrying capacity (*K* = 6) through another transcritical bifurcation visible in the *P* − *F* portrait, a stable equilibrium is obtained with the predator feeding on the healthy and infected prey populations.

The phase portraits reveal how for a low prey carrying capacity only the healthy prey can survive in the system (Figure 5A), but at increasing carrying capacities, a first transcritical bifurcation makes the infected prey population to be able to coexist with the healthy prey (Figure 5B). A second transcritical bifurcation results in the three populations to survive stably in the system (Figure 5C). The portraits clearly show how the nullclines cross when increasing the carrying capacity to displace the stable critical point to non-zero values for the infected population first and then for the predator population as well.

More complex dynamics occur in this three-dimensional model at higher prey carrying capacities. Figure 6 shows how the generated portraits help understand the emergent dynamics. As the system gets closer to a subcritical Hopf bifurcation, the amplitude of the oscillations towards the stable coexisting attractor increases (Figure 6A). However, after crossing the bifurcation, the oscillations get critically close to zero population values, which causes the system to collapse, revealing an unstable limit cycle (Figure 6B). Further increasing the prey carrying capacity produces an earlier collapse of the system (Figure 6C). The portraits illustrate how these collapse dynamics occur as the initially stable critical point gets closer to the zero axes, which finally results in the populations to asymptotically reach extinction. Interacting with the portraits through the interface by varying the values of the parameters, the viewed planes, and the trajectories can facilitate the understanding of these complex dynamics.

**Figure 6.**
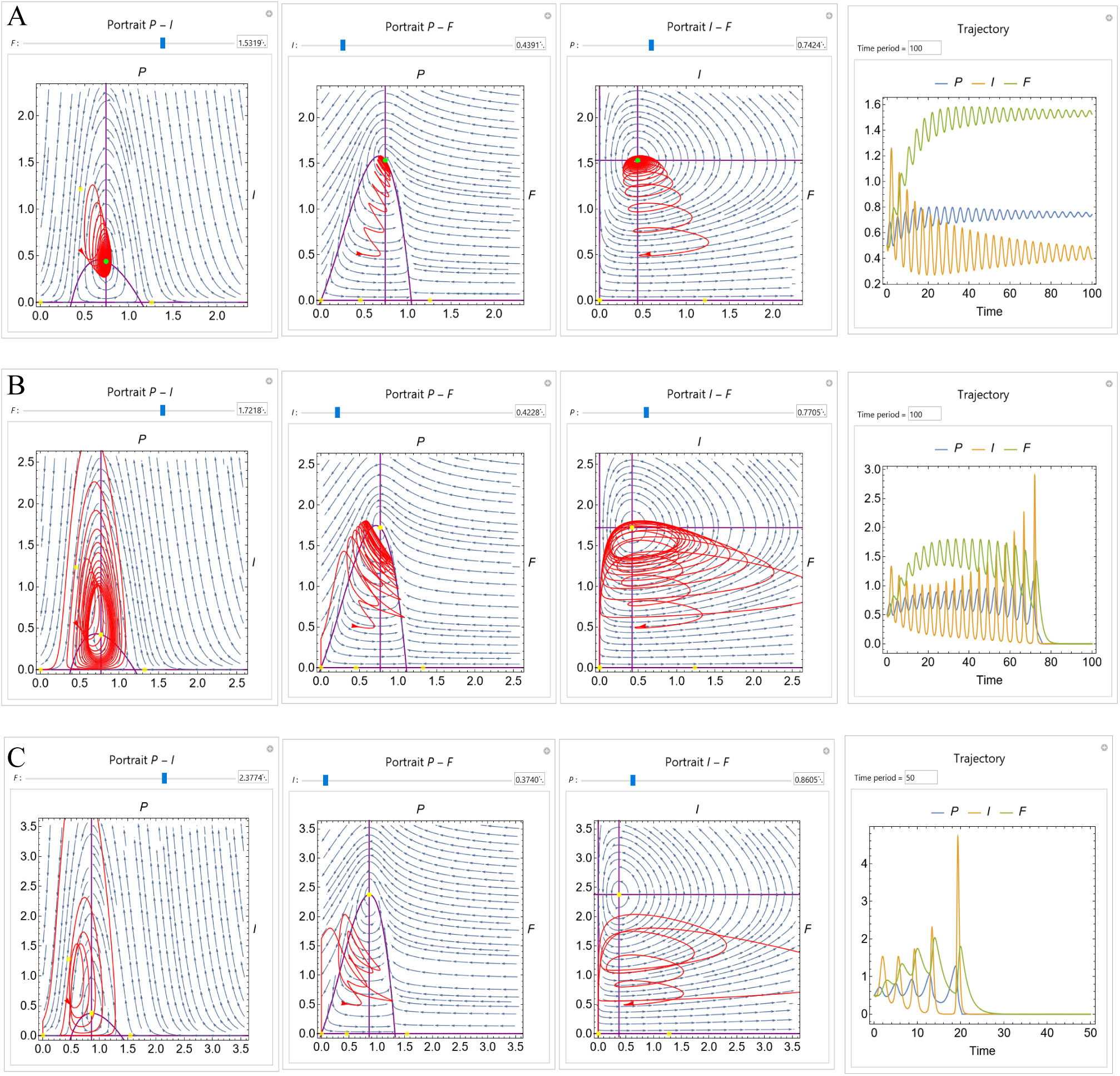
Outputs of the method for the predator (*F*), prey (*P*) model including prey group defense and an infected prey population (*P*) with high levels of prey carrying capacity *K*. All other parameters are set as in Table 1. All the portraits are automatically centered on the endemic equilibrium point and show a trajectory (red and right panels) starting at a population of 0.5 for all the species. **A**. When increasing the prey carrying capacity (*K* = 9.9), the endemic predator-prey equilibrium is still a stable attractor (green dot), albeit with higher oscillations (see Fig. 5C). **B**. After increasing the prey carrying capacity (*K* = 11) through a subcritical Hopf bifurcation, the endemic predator-prey equilibrium becomes unstable, and an unstable limit cycle appears after which the system collapses. **C**. Further increasing the prey carrying capacity (*K* = 15) shortens the time for the system to collapse, as the unstable equilibrium gets closer to the asymptotic dynamics at the horizontal and vertical axes shown in the portraits.

### 4.2. Lotka-Volterra competition model

Lotka-Volterra competition models with four or more species can exhibit chaotic behaviors [22]. We consider the four-species model studied in [23], defined for populations *x*_*i*_, *i* = (1, …, 4) and governed with dynamics

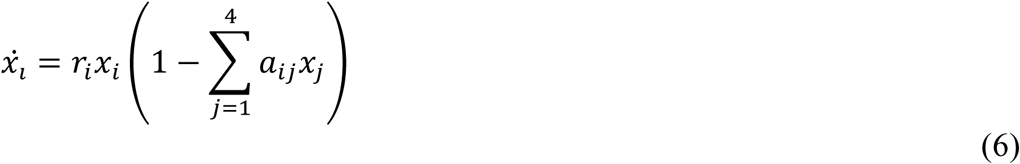

where *r*_*i*_ is the intrinsic growth rate of species *i*, and *a*_*ij*_ is the species *j* competition factor for resources used by species *i*. A set of parameters that produce chaotic solutions in the system were previously found using a numerical approach based on simulated annealing [23], resulting in parameter values:

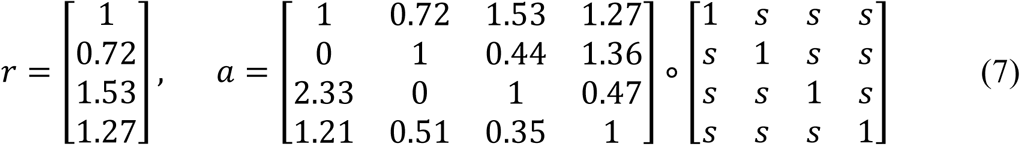

where the free parameter *s* multiplies the off-diagonal elements of *a*, modulating the coupling strength between species. Here we study the dynamics of the system with automatically-generated phase portraits for different degrees of coupling strength.

When the coupling strength parameter is set to *s* = 0, the system of equations become uncoupled and each population follows an independent Verhulst logistic growth model. Figure 7 shows the generated portraits with the presented method for this case. The four populations have the same stable attractor at a value of 1. Activating the coupling between the four populations (*s* = 0.8) results in the equilibrium point for all the populations shifting towards zero. This shift causes the dynamics to oscillate in the form of a spiral towards the coexistence stable attractor (Figure 8). Increasing the coupling parameter (*s* = 0.9) causes the system to pass through a Hopf bifurcation, resulting in the coexistence attractor becoming unstable (Figure 9). In this case, the populations do not reach a single value equilibrium but oscillate indefinitely towards a limit cycle. Further increasing the interdependency (*s* = 0.96) doubles the period of the oscillations with a harmonic of the dominant frequency (Figure 10). The doubled period is visible in the plotted trajectory with the two thicker rings. Finally, setting the parameters to the values found with the heuristic search (s=1), reveals the chaotic attractor (Figure 11). Although the trajectories pass close the asymptotic zero axis for population *x*_3_, they always move back around the strange attractor now formed by the unstable coexistence equilibrium point.

**Figure 7.**
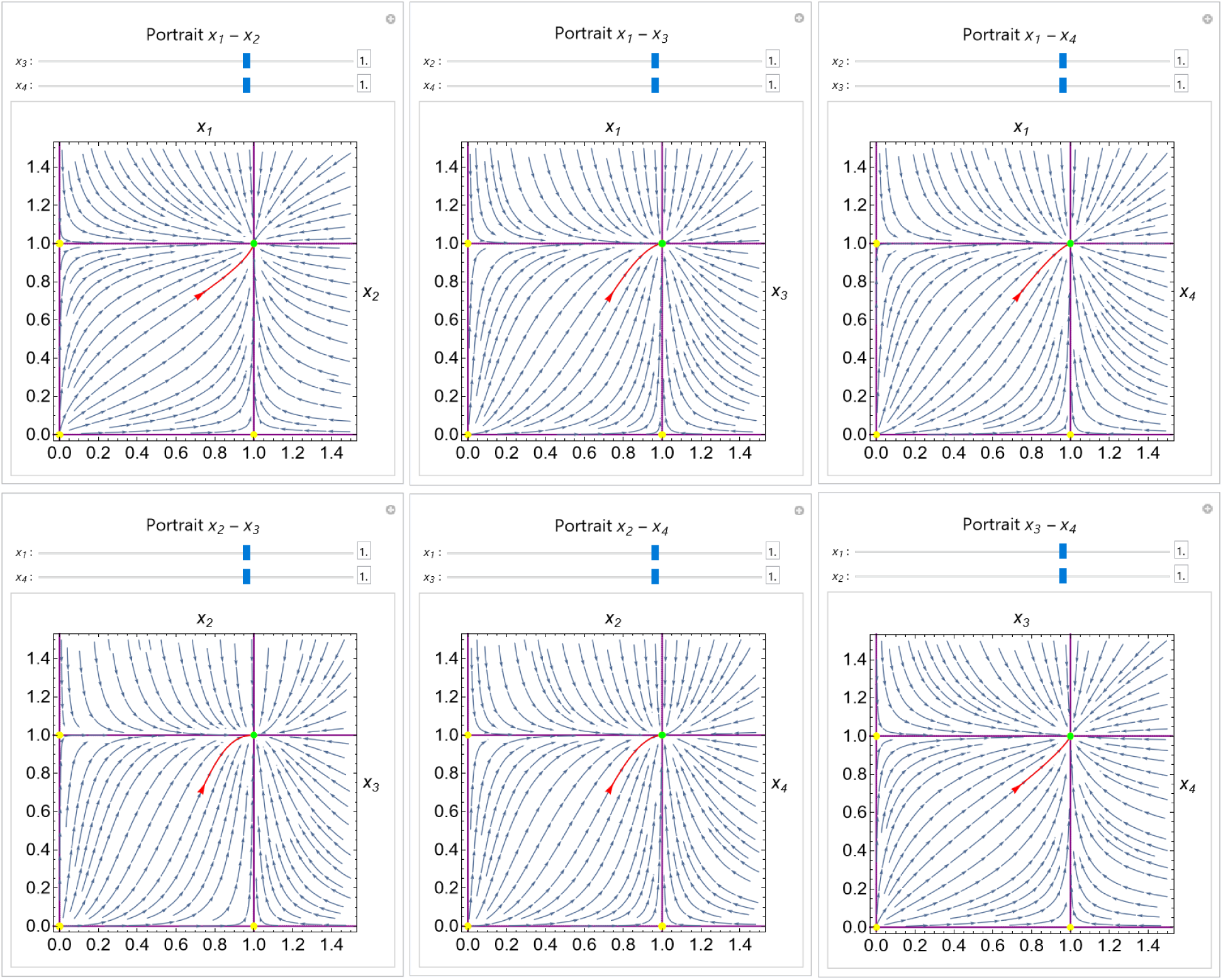
Phase portraits of a four-species Lotka Volterra model of competition without coupling (*s* = 0). The portraits are automatically centered at the plane containing the stable critical point. A trajectory (red line) shows the attractor dynamics towards coexistence equilibrium.

**Figure 8.**
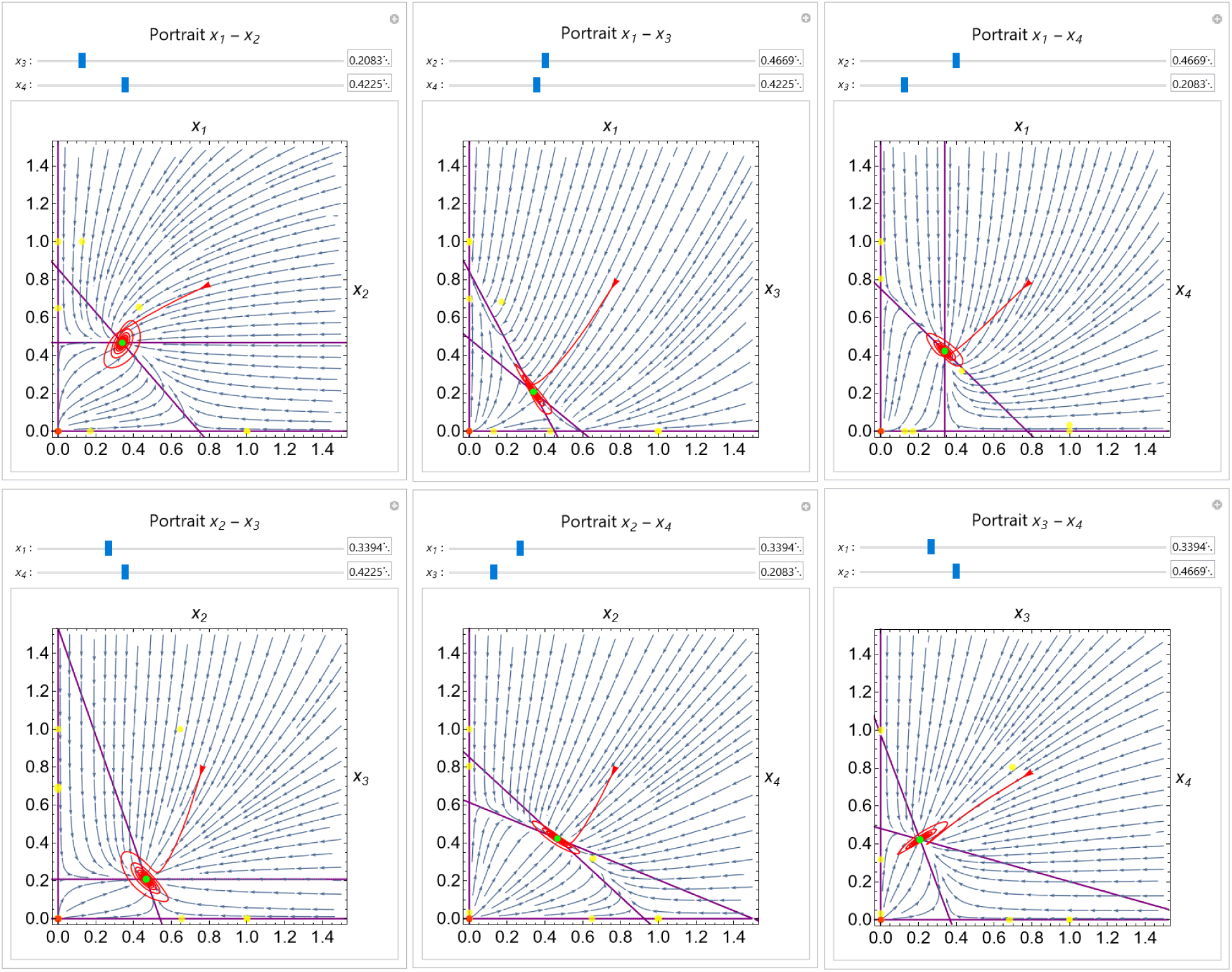
Phase portraits of a four-species Lotka Volterra model of competition with low coupling (*s* = 0.8). A trajectory (red) shows how the dynamics of the system present spiral oscillations towards the coexistence equilibrium state (green dot).

**Figure 9.**
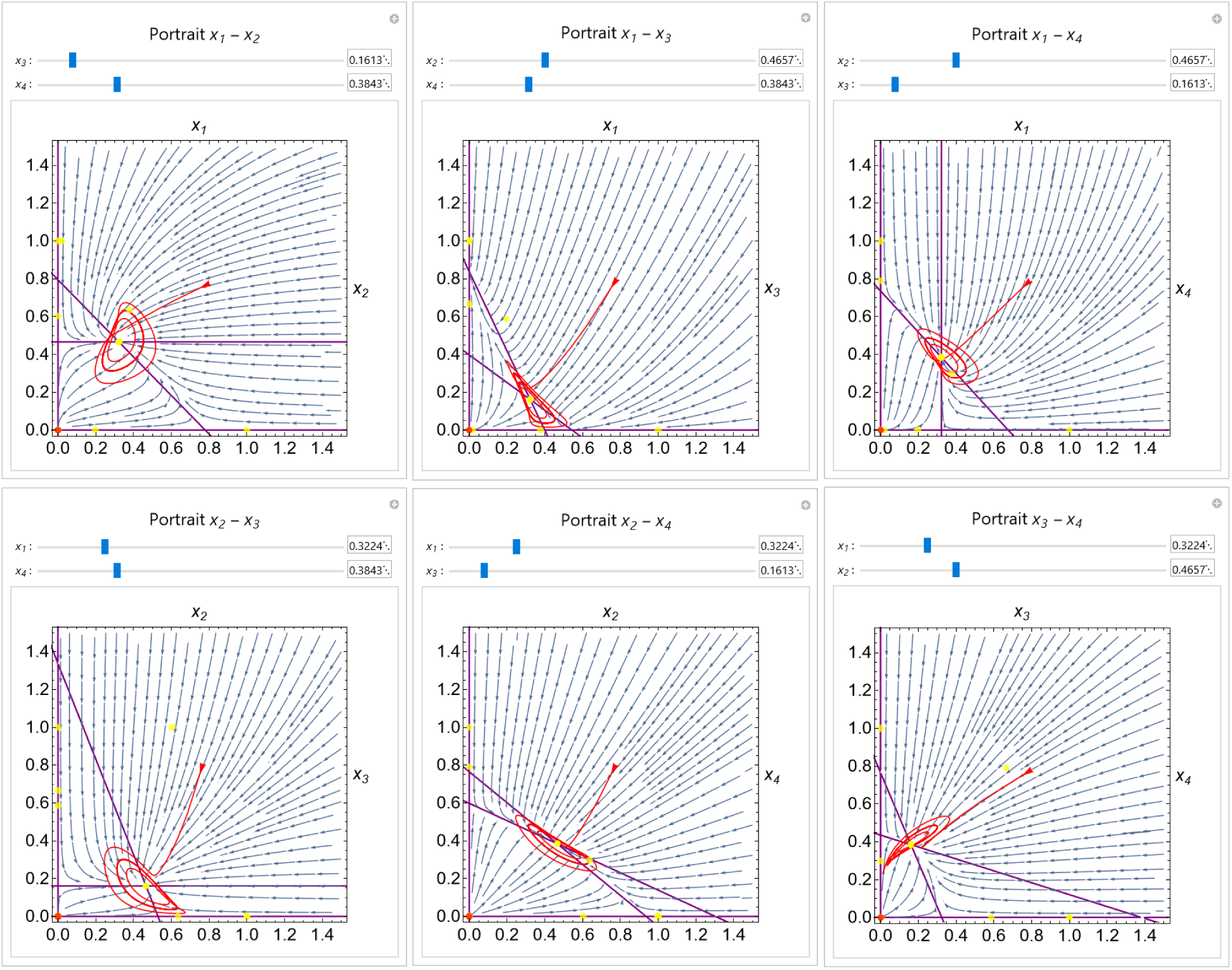
Phase portraits of a four-species Lotka Volterra model of competition with increased coupling (*s* = 0.9). A Hopf bifurcation causes the coexistence equilibrium to become unstable, and the system instead spirals towards a limit cycle, as shown by the highlighted trajectory (red).

**Figure 10.**
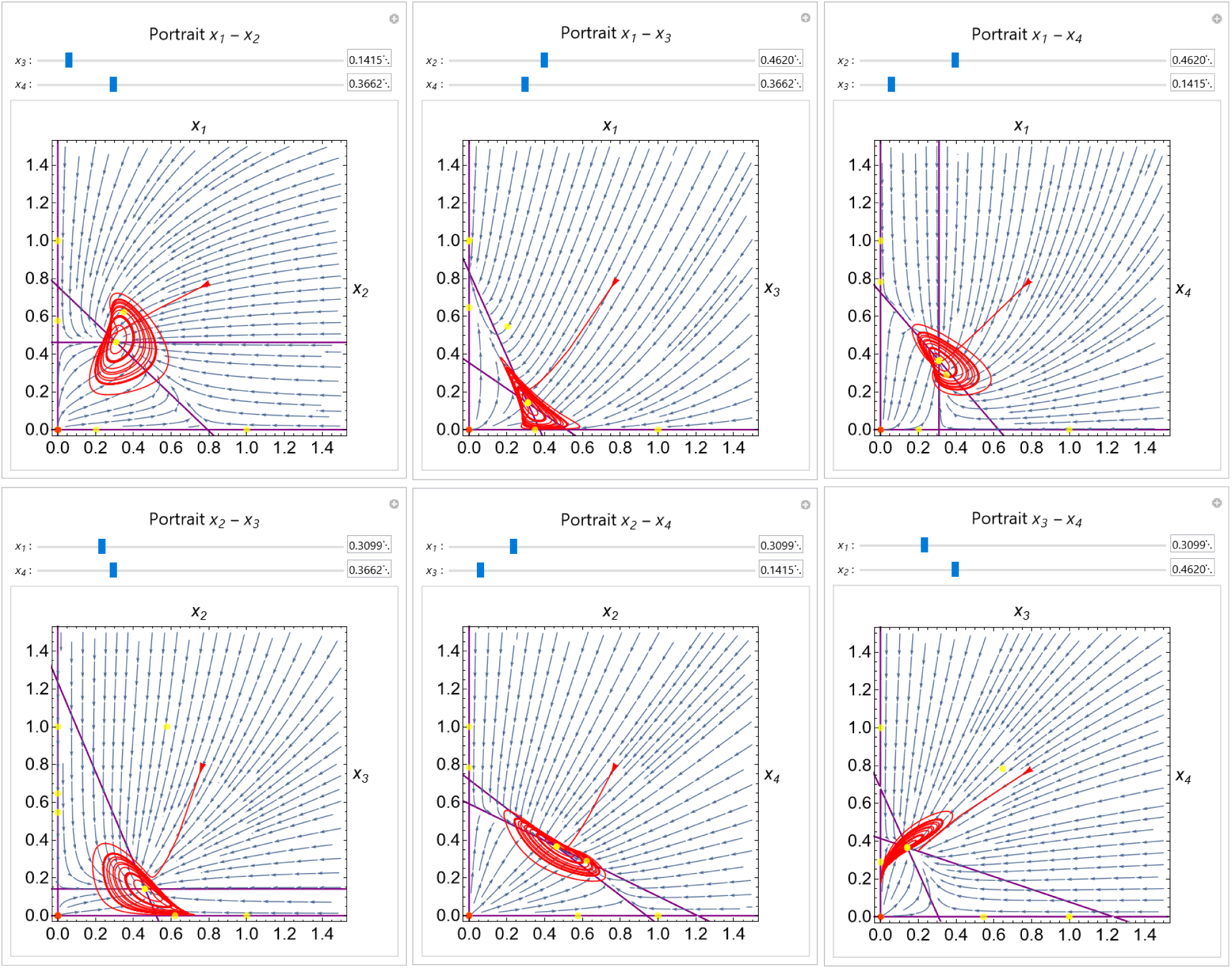
Phase portraits of a four-species Lotka Volterra model of competition with high coupling (*s* = 0.96). The oscillations period is doubled, as revealed with the two thicker lines displayed by the trajectory (red), especially visible in the *x*_1_ − *x*_2_ plot.

**Figure 11.**
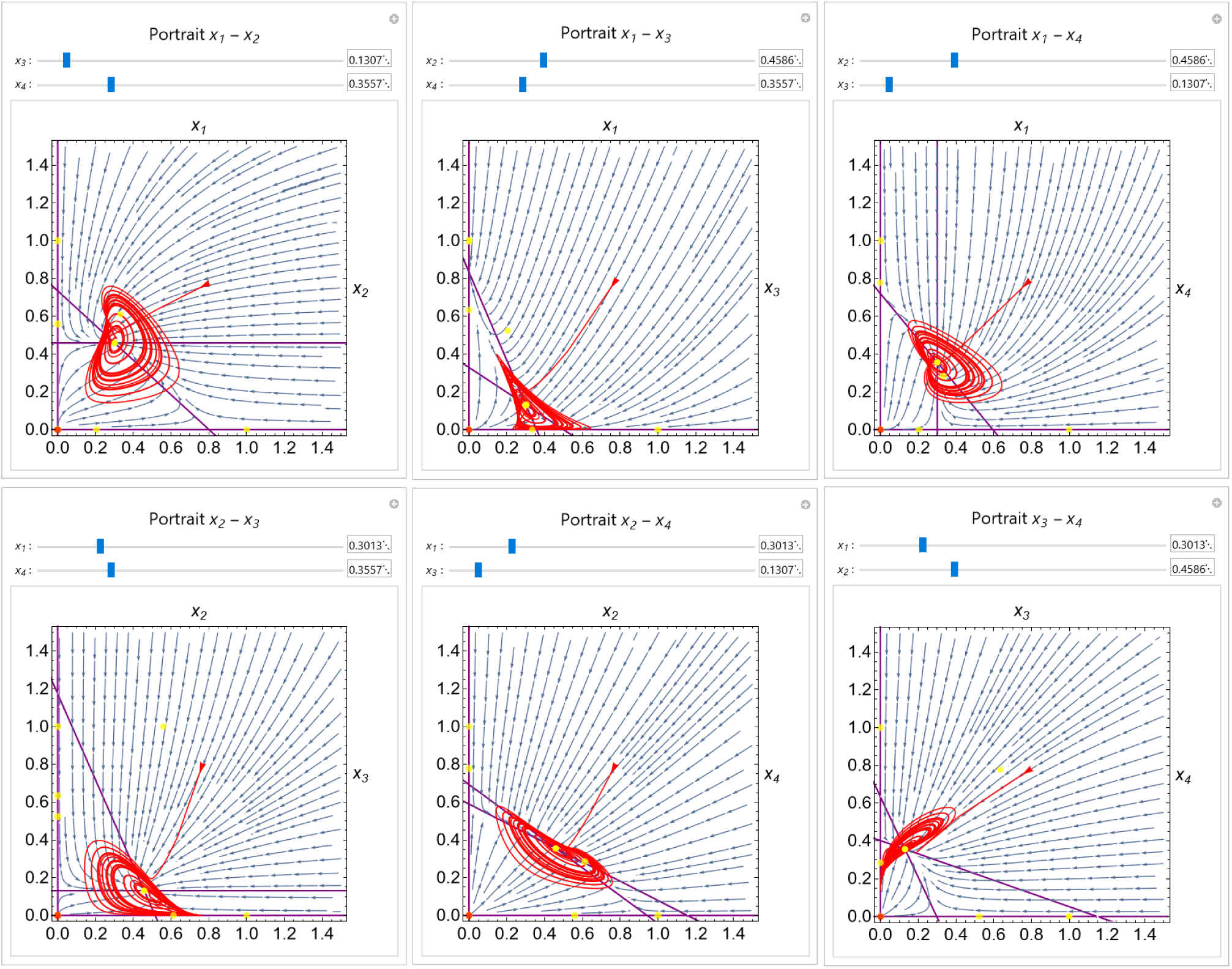
Phase portraits of a four-species Lotka Volterra model of competition displaying a chaotic behavior (*s* = 1).

The trajectories for the four-species Lotka Volterra model of competition at different coupling strengths are shown in the time domain in Figure 12 as generated by the presented method. The plots make clear how as the coupling strength increases, the dynamics of the system become more complex, from equilibrium attractors, to limit cycles, and finally a chaotic behavior. Indeed, the trajectory visualizations in the time domain complement the phase portraits to facilitate the understanding of these complex dynamics.

**Figure 12.**
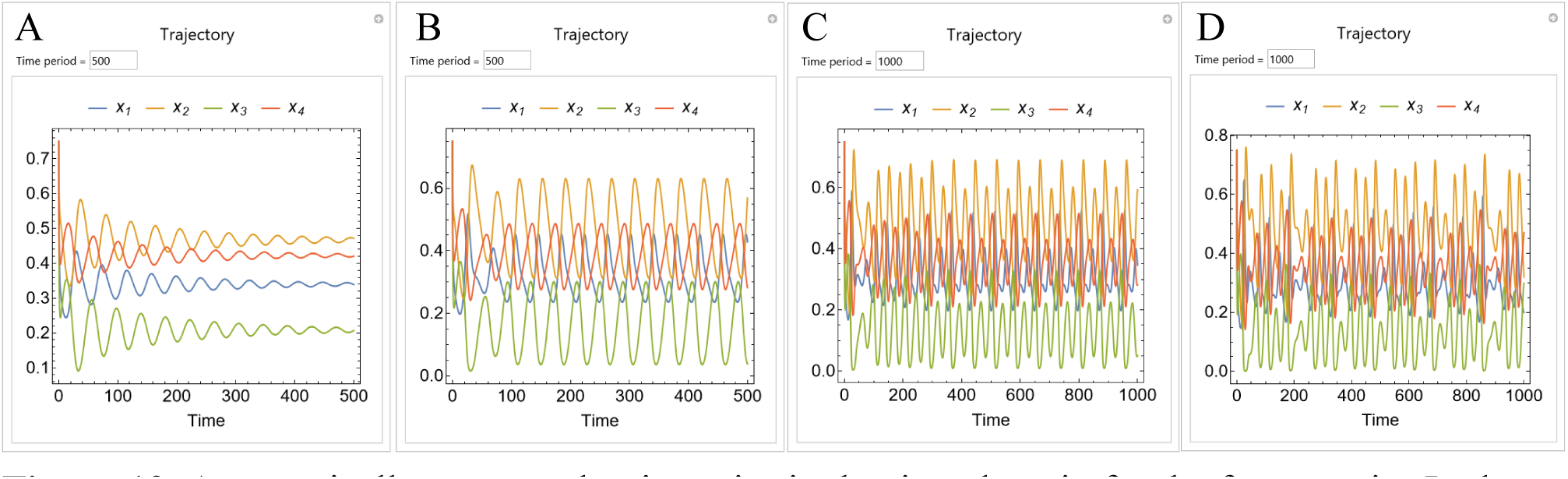
Automatically-generated trajectories in the time domain for the four-species Lotka Volterra model of competition at different coupling strengths. Each trajectory shown in panels A-D corresponds to the trajectory shown in Figures 8-11, respectively, and start at value 0.75 for all the species. **A**. At low coupling strength (*s* = 0.8), the system converges to a coexistence equilibrium. **B**. Increasing the coupling strength (*s* = 0.9) through a Hopf bifurcation, the system becomes cyclic. **C**. A further increase of the coupling strength (*s* = 0.96) doubles the oscillation period. **D**. At complete coupling strength (*s* = 1), the system becomes chaotic.

## 5. Discussion and conclusion

We have presented a new method called *AutoPortrait* to automatically generate interactive phase portraits of biological dynamical systems. The method can take as input multidimensional systems of differential equations and produce a phase portrait for each pair of dependent variables in order to understand their complex dynamics. The portraits include a vector field, nullclines, and the critical points of the system. In addition, the phase portraits can include trajectories that are displayed as projections in all the planes shown in the portraits as well as in a visualization in the time domain. Altogether, these portraits allow for the complete understanding of the dynamics of a complex biological system.

Crucially, the automatically-generated phase portraits are interactive. The user can pan and zoom the plots using the mouse as well as center the viewed plane clicking the different critical points. For systems with more than two dimensions, sliders and input boxes for each of the dependent variables not included in each of the phase portraits can be used to change the plane visualized. The parameters in the system are automatically recognized by the method, and the user can change their values with sliders or input boxes. All these available user interactions result in the update of the phase portrait in real time, which facilitates the exploration and comprehension of the system.

The generated phase portraits can be copied and integrated into Mathematica notebooks retaining all the interactive functionality, an ideal solution for dynamic publication media and pedagogical materials. These interactive notebooks can be opened with the freely-available *Wolfram Player* in the most commonly-used operating systems, facilitating the diffusion and accessibility of the produced materials. Importantly, such dynamic computational notebooks can be adopted as supplementary dynamic media, a crucial step towards reproducible research in mathematical sciences [24].

In order to automatically generate the phase portraits with the presented method, we have implemented a *Mathematica* package that can be imported into scripts, in addition to a user-friendly graphical user interface. The graphical interface allows any user without programming knowledge to directly type the system of equations into input boxes and click a button to automatically generate their portraits. This represents an excellent method for the fast generation of portraits as well as a valuable educational tool to be used in an interactive classroom for dynamical systems or systems biology [25]. The package interface allows the generation of interactive portraits programmatically and the integration of the methodology in other computational pipelines. This approach could be especially useful in methodologies that can infer the parameters or the entire dynamic model from phenotypic data [26–28], facilitating the understanding of these automatically-generated mathematical models.

In conclusion, the presented method represents a novel tool for the automatic generation of phase portraits for multidimensional biological systems. The ability to interact in real time with the portraits, as well as to save a specific system and particular views in a computable notebook makes it especially useful to disseminate reproducible research results and interactive pedagogical materials. Furthermore, the provided user-friendly interface can be used without any programming knowledge, which is a valuable asset as an educational tool in the classroom [29] and for promoting reproducible research in biology modeling [30].

## Acknowledgments

We thank the members of the Lobo Lab for helpful discussions. This work was supported by the National Institute of General Medical Sciences of the National Institutes of Health under award number R35GM137953; O.O.O. was supported by the U-RISE program at UMBC under award number T34GM136497. The content is solely the responsibility of the authors and does not necessarily represent the official views of the National Institutes of Health.

## References

[1] J.M. Ko, R. Mousavi, D. Lobo, Computational Systems Biology of Morphogenesis, in: S. Cortassa, M. Aon (Eds.), Computational Systems Biology in Medicine and Biotechnology, Springer US, 2022: pp. 343–365. https://doi.org/10.1007/978-1-0716-1831-8_14.

[2] Z. Ji, K. Yan, W. Li, H. Hu, X. Zhu, Mathematical and Computational Modeling in Complex Biological Systems, BioMed Research International. 2017 (2017) 1–16. https://doi.org/10.1155/2017/5958321.

[3] S. Hoops, R. Gauges, C. Lee, J. Pahle, N. Simus, M. Singhal, L. Xu, P. Mendes, U. Kummer, COPASI - A COmplex PAthway SImulator, Bioinformatics. 22 (2006) 3067–3074. https://doi.org/10.1093/bioinformatics/btl485.

[4] B. Shaikh, G. Marupilla, M. Wilson, M.L. Blinov, I.I. Moraru, J.R. Karr, RunBioSimulations: An extensible web application that simulates a wide range of computational modeling frameworks, algorithms, and formats, Nucleic Acids Research. 49 (2021) W597–W602. https://doi.org/10.1093/nar/gkab411.

[5] R.S. Malik-Sheriff, M. Glont, T.V.N. Nguyen, K. Tiwari, M.G. Roberts, A. Xavier, M.T. Vu, J. Men, M. Maire, S. Kananathan, E.L. Fairbanks, J.P. Meyer, C. Arankalle, T.M. Varusai, V. Knight-Schrijver, L. Li, C. Dueñas-Roca, G. Dass, S.M. Keating, Y.M. Park, N. Buso, N. Rodriguez, M. Hucka, H. Hermjakob, BioModels—15 years of sharing computational models in life science, Nucleic Acids Research. 48 (2019) D407–D415. https://doi.org/10.1093/nar/gkz1055.

[6] S. Ghosh, Y. Matsuoka, Y. Asai, K. Hsin, H. Kitano, Software for systems biology : from tools to integrated platforms, Nature Publishing Group. 12 (2011) 821–832. https://doi.org/10.1038/nrg3096.

[7] T. Helikar, B. Kowal, S. McClenathan, M. Bruckner, T. Rowley, A. Madrahimov, Wicks, M. Shrestha, K. Limbu, J. Rogers, The Cell Collective: Toward an open and collaborative approach to systems biology, BMC Systems Biology. 6 (2012) 96.

[8] T. Helikar, The Need for Research-Grade Systems Modeling Technologies for Life Science Education, Trends in Molecular Medicine. 27 (2021) 100–103. https://doi.org/10.1016/j.molmed.2020.11.005.

[9] J. Jaeger, N. Monk, Bioattractors: dynamical systems theory and the evolution of regulatory processes, The Journal of Physiology. 592 (2014) 2267–2281. https://doi.org/10.1113/jphysiol.2014.272385.

[10] S. Green, M. Fagan, J. Jaeger, Explanatory Integration Challenges in Evolutionary Systems Biology, Biological Theory. 10 (2015) 18–35. https://doi.org/10.1007/s13752-014-0185-8.

[11] S.H. Strogatz, Nonlinear Dynamics and Chaos, 2nd editio, CRC Press, 2015. https://www.taylorfrancis.com/books/9780429961113.

[12] M.A. Rodríguez-Licea, F.J. Perez-Pinal, J.C. Nuñez-Pérez, Y. Sandoval-Ibarra, On the n-dimensional phase portraits, Applied Sciences (Switzerland). 9 (2019) 1–19. https://doi.org/10.3390/app9050872.

[13] J. Musilova, K. Sedlar, Tools for time-course simulation in systems biology: A brief overview, Briefings in Bioinformatics. 22 (2021) 1–15. https://doi.org/10.1093/bib/bbaa392.

[14] R.W. Shonkwiler, J. Herod, Mathematical Biology: An Introduction with Maple and Matlab, Springer New York, New York, NY, 2009. https://doi.org/10.1007/978-0-387-70984-0.

[15] S. Lynch, Dynamical Systems with Applications using Mathematica®, Birkhäuser Boston, Boston, MA, 2007. https://doi.org/10.1007/978-0-8176-4586-1.

[16] B. Ermentrout, Simulating, Analyzing, and Animating Dynamical Systems: A Guide to XPPAUT for Researchers and Students, Society for Industrial and Applied Mathematics, 2002. https://doi.org/10.1137/1.9780898718195.

[17] T. Müller, F. Sadlo, Visual exploration of 2D autonomous dynamical systems, European Journal of Physics. 36 (2015) 1–11. https://doi.org/10.1088/0143-0807/36/3/035007.

[18] I. Ahmad, A. Saaban, A. Ibrahim, M. Shahzad, A Research on Active Control to Synchronize a New 3D Chaotic System, Systems. 4 (2015) 2. https://doi.org/10.3390/systems4010002.

[19] G. Ge, W. Wang, The Application of the Undetermined Fundamental Frequency Method on the Period-Doubling Bifurcation of the 3D Nonlinear System, Abstract and Applied Analysis. 2013 (2013) 1–6. https://doi.org/10.1155/2013/813957.

[20] E.O. Voit, A First Course in Systems Biology, 2nd ed., Garland Science, Second edition. | New York : Garland Science, 2017., 2017. https://doi.org/10.4324/9780203702260.

[21] B.W. Kooi, E. Venturino, Ecoepidemic predator-prey model with feeding satiation, prey herd behavior and abandoned infected prey, Mathematical Biosciences. 274 (2016) 58–72. https://doi.org/10.1016/j.mbs.2016.02.003.

[22] R. Wang, D. Xiao, Bifurcations and chaotic dynamics in a 4-dimensional competitive Lotka-Volterra system, Nonlinear Dynamics. 59 (2010) 411–422. https://doi.org/10.1007/s11071-009-9547-3.

[23] J.A. Vano, J.C. Wildenberg, M.B. Anderson, J.K. Noel, J.C. Sprott, Chaos in low-dimensional Lotka-Volterra models of competition, Nonlinearity. 19 (2006) 2391–2404. https://doi.org/10.1088/0951-7715/19/10/006.

[24] S. Schnell, “Reproducible” Research in Mathematical Sciences Requires Changes in our Peer Review Culture and Modernization of our Current Publication Approach, Bulletin of Mathematical Biology. 80 (2018) 3095–3105. https://doi.org/10.1007/s11538-018-0500-9.

[25] T. Helikar, C.E. Cutucache, L.M. Dahlquist, T.A. Herek, J.J. Larson, J.A. Rogers, Integrating Interactive Computational Modeling in Biology Curricula, PLoS Comput Biol. 11 (2015) e1004131. https://doi.org/10.1371/journal.pcbi.1004131.

[26] R. Mousavi, S.H. Konuru, D. Lobo, Inference of dynamic spatial GRN models with multi-GPU evolutionary computation, Briefings in Bioinformatics. 22 (2021) 1–11. https://doi.org/10.1093/bib/bbab104.

[27] J. Hwang, A. Hari, R. Cheng, J.G. Gardner, D. Lobo, Kinetic modeling of microbial growth, enzyme activity, and gene deletions: An integrated model of β-glucosidase function in Cellvibrio japonicus, Biotechnology and Bioengineering. 117 (2020) 3876–3890. https://doi.org/10.1002/bit.27544.

[28] D. Lobo, Formalizing Phenotypes of Regeneration, in: Whole-Body Regeneration, Springer US, 2022. https://doi.org/10.1007/978-1-0716-2172-1_36.

[29] M. Evagorou, K. Korfiatis, C. Nicolaou, C. Constantinou, An investigation of the potential of interactive simulations for developing system thinking skills in elementary school: A case study with fifth-graders and sixth-graders, International Journal of Science Education. 31 (2009) 655–674. https://doi.org/10.1080/09500690701749313.

[30] K. Tiwari, S. Kananathan, M.G. Roberts, J.P. Meyer, M.U. Sharif Shohan, A. Xavier, M. Maire, A. Zyoud, J. Men, S. Ng, T.V.N. Nguyen, M. Glont, H. Hermjakob, R.S. Malik-Sheriff, Reproducibility in systems biology modelling, Molecular Systems Biology. 17 (2021) 1–7. https://doi.org/10.15252/msb.20209982.

